# Intercellular viral spread and intracellular transposition of *Drosophila* gypsy

**DOI:** 10.1101/2020.05.28.121897

**Authors:** Richard M. Keegan, Yung-Heng Chang, Michael J. Metzger, Josh Dubnau

**Affiliations:** Program in Neuroscience, Department of Neurobiology and Behavior, Stony Brook University, NY 11794, USA; Department of Anesthesiology, Stony Brook School of Medicine, NY 11794, USA; Pacific Northwest Research Institute, Seattle, WA 98122, USA

## Abstract

It has become increasingly clear that retrotransposons (RTEs) are more widely expressed in somatic tissues than previously appreciated. RTE expression has been implicated in a myriad of biological processes ranging from normal development and aging, to age related diseases such as cancer and neurodegeneration. Long Terminal Repeat (LTR)-retrotransposons are evolutionary ancestors to, and share many features with, exogenous retroviruses. In fact, many organisms contain endogenous retroviruses (ERVs) that derive from an exogenous retrovirus that have integrated into the germ line. These ERVs are inherited in Mendelian fashion like RTEs, and some retain the ability to transmit between cells like viruses, while others develop the ability to act as RTEs. The process of evolutionary transition between LTR-RTE and retroviruses is thought to involve multiple steps by which the element loses or gains the ability to transmit copies between cells versus the ability to replicate intracellularly. But, typically, these two modes of transmission are incompatible because they require assembly in different sub-cellular compartments. Like murine IAP/IAP-E elements, the gypsy family of retroelements in arthropods appear to sit along this evolutionary transition. The fact that gypsy elements have been found to actively mobilize in neurons and glial cells during normal aging and in models of neurodegeneration raises the question of whether their replication in somatic cells occurs via intracellular retrotransposition, intercellular viral spread, or some combination of the two. These modes of replication in somatic tissues would have quite different biological implications. Here, we demonstrate that Drosophila gypsy is capable of both cell-associated and cell-free viral transmission between cultured S2 cells of somatic origin. Further, we demonstrate that the ability of gypsy to move between cells is dependent upon a functional copy of its viral envelope protein. This argues that the gypsy element has transitioned from an RTE into a functional endogenous retrovirus with the acquisition of its envelope gene. On the other hand, we also find that intracellular retrotransposition of the same genomic copy of gypsy can occur in the absence of the Env protein. Thus, gypsy exhibits both intracellular retrotransposition and intercellular viral transmission as modes of replicating its genome.

## Introduction

The genomes of plants and animals contain a substantial contribution of sequences derived from transposable elements (TEs). In humans, for example, TE derived sequences represent nearly half of all genetic material (1). TEs mainly act as selfish genetic elements that replicate within germline tissue, where their de novo inserted copies can be passed to offspring, allowing vertical spread within a population (2, 3). But in the case of the Type I TEs, known as retrotransposons (RTEs), there is now compelling evidence that expression and even replication also occurs in somatic tissues and impacts both normal biology and a variety of age-related diseases (4-7).

Members of the LINE, LTR-RTE, and ERV families of RTEs have been found to be actively expressed and even to replicate in somatic tissues, most notably within the nervous system (4, 6-25). Although functional consequences of RTE replication during normal neural development are not established, there is growing evidence that dysfunctional expression has a detrimental impact on organismal fitness during aging (11, 16, 26-37) and in age-related diseases such as cancer (38-52), autoimmune disorders (53-55) and neurodegenerative disorders such as amyotrophic lateral sclerosis (10, 11, 56-61), frontotemporal dementia (59), Aicardi-Goutieres syndrome (62, 63), Alzheimer’s (64-68), progressive supranuclear palsy (67), multiple sclerosis (69-71), fragile X-associated tremor/ataxia syndrome (72), macular degeneration (73), and Rett syndrome (74).

Like retroviruses, RTEs replicate through an RNA intermediate which is then converted into DNA by an encoded reverse transcriptase enzyme. DNA copies can be inserted into de novo chromosomal sites in the genome, thereby increasing copy number with each successive replication cycle (75-77). Indeed, a subset of RTEs, the so-called long terminal repeat (LTR)-RTEs, are evolutionarily related to retroviruses. Unlike exogenous retroviruses, both long interspersed nuclear element (LINE) and LTR-RTEs are primarily adapted to make use of an intracellular replication cycle, although there is some evidence for transfer via extracellular vesicles (78-80). Functional LTR-RTEs encode gag and pol open reading frames, but unlike retroviruses they do not contain an envelope ‘glycoprotein (Env) to mediate inter-cellular spread. Also, they generally target assembly of virus-like particles at the lumen of the ER to facilitate re-entry to the nucleus rather than at the extracellular membrane to facilitate release from the cell (81).

Such LTR-RTEs are believed to be the evolutionary ancestors of exogenous retroviruses, which emerged by a multi-step process that includes the gain of an *Env* gene (82-85) and re-targeting of assembly to the extracellular membrane. This process also has occurred in reverse, leading to endogenous retroviruses (ERVs) that over time can lose their *Env* gene and re-target their assembly for intracellular replication, acting like LTR-RTEs. Indeed, many genomes contain such ERVs, which straddle the evolutionary transition between LTR-retrotransposon and exogenous retrovirus. Gypsy elements in *Drosophila*, the murine IAP-E elements and the HERV-K elements in human genomes, for example, each retain the viral *Env*, and may therefore have the potential to act as either a virus or a retrotransposon.

Although the bulk of research into somatic retrotransposition has so far focused on LINE elements (5), the gypsy ERV also has been shown capable of replicating in somatic tissues in *Drosophila*, including glial cells, post-mitotic neurons, adipose tissues, and intestinal stem cells (10, 11, 16, 33, 58, 67, 86, 87), and HERV-K expression has been detected in ALS patients and in several cancers (57, 60, 61, 88-90). The expression and replication within somatic tissues of ERVs, which encode functional Env proteins, highlights the importance of understanding their replication cycle. These elements sit on a spectrum between intracellular RTE and extracellular virus. It is not clear whether such elements replicate through intracellular transposition or whether their replication requires them to move genetic material between somatic cells via viral transmission (81, 91-97)

We have addressed this question using cultured *Drosophila* S2 cells of macrophage lineage. We used a replication reporter system that we recently developed (11) as well as a series of novel reporters, to test whether or not gypsy replication occurs via intra-cellular transposition or intercellular viral transfer. We find that gypsy can transfer between separate populations of cells in cell culture using both cell-free and cell-associated modes of transmission. We further demonstrate that both forms of transmission between cells requires an intact Env open reading frame (ORF). Surprisingly, we also find that in the absence of Env, gypsy is able to efficiently complete intracellular retrotransposition.

## Results

### GYPSY-CLEVR AND GYPSY-MCHERRY REPORTERS OF GYPSY REPLICATION AND EXPRESSION

We previously described a gypsy reporter system, **C**ellular **L**abeling of **E**ndogenous Retro**v**irus **R**eplication (CLEVR). The gypsy-CLEVR reporter reliably marks cells in which replication of gypsy has occurred and in which a de novo cDNA copy has been reinserted into the genome. This reporter system reliably reports replication of the exogenously supplied gypsy construct both in cell culture and in vivo (10, 11). This gypsy-CLEVR reporter contains the full-length gypsy sequence with a promoterless watermelon (WM) dual fluorescent gene in the 3’LTR and a Gal4-sensitive promoter in the 5’LTR, and it takes advantage of the conserved template switching steps in retrovirus replication to place the Gal4-sensitive promoter upstream to the WM reporter. The gypsy-CLEVR reporter expression requires the replication of gypsy to link the promoter to the reporter and requires the presence of Gal4 to drive the WM signal after replication (11). The gypsy-CLEVR reporter, and control versions that are unable to replicate due to mutations in the essential primer binding site (PBS) were employed here (11) (Figure 1A). To examine inter-cellular spread of gypsy, we also generated gypsy-mCherry, a more standard reporter of gypsy expression. Gypsy-mCherry relies on the porcine teschovirus-1 2A (P2A) self-cleaving peptide (98) to link expression of mCherry to expression of *Env* (Figure 1A). In contrast with the gypsy-CLEVR reporter, gypsy-mCherry marks any cells in which the cconstruct is expressed, differing from CLEVR in that it does not require replication. As the translation of mCherry is linked directly to the env encoding (spliced) transcript of gypsy, this reporter is driven by the gypsy-endogenous promoter and does not require Gal4 to display fluorescent signal. We also generated a version of this construct in which the Env protein coding sequence was deleted (Figure 1A).

**Figure 1:**
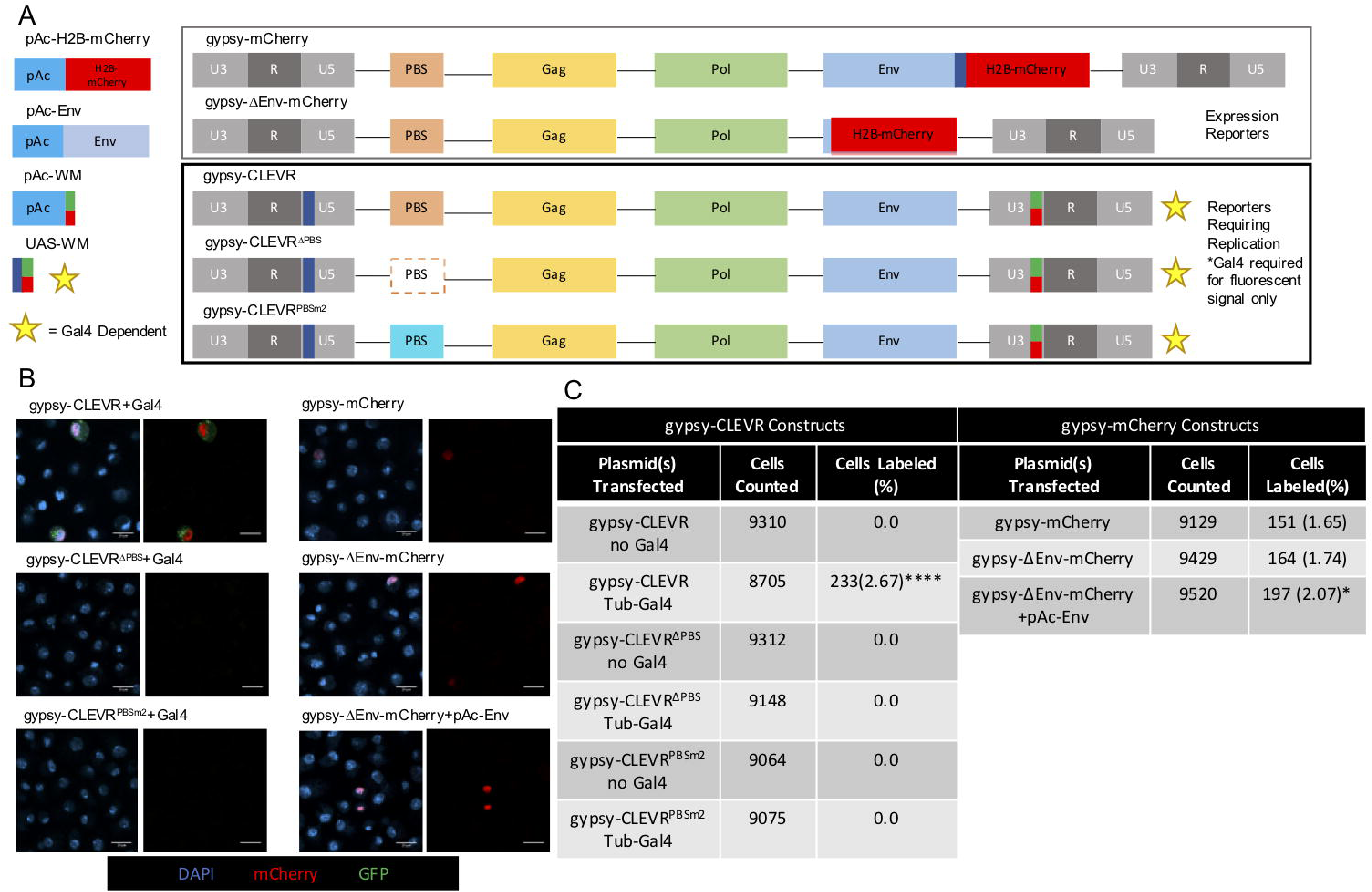
Functional test of reporters marking gypsy replication and expression in Drosophila S2 cells. (A) Cartoon representations of the pAc-H2B-mCherry, pAc-Env, pAc-WM, UAS-WM, gypsy-CLEVR, gypsy-CLEVR^ΔPBS^, gypsy-CLEVR^PBSm2^, gypsy-mCherry, and gypsy-ΔEnv-mCherry constructs used. Star denotes Gal4 dependence for fluorescence but not replication. (B) Fluorescent images showing WM or mCherry positive S2 cells for gypsy-CLEVR and gypsy-mCherry constructs. Scale bars = 10 µm. (C) Quantification showing the percentage of cells labeled for the gypsy-CLEVR constructs with and without Gal4, and the gypsy-mCherry constructs. Quantification is presented as totals cells counted from 3 near equivalent sets of biological replicates. Significance for the gypsy-CLEVR constructs was calculated against gypsy-CLEVR with no Gal4; significance for gypsy-mCherry constructs was calculated against gypsy-mCherry. Significance was determined using the Fisher’s Exact test variant of the Chi^2^ test. Significance values are denoted as: p=<0.05 *, p=<0.001 ***, p=<0.0001****

To confirm the fidelity of the gypsy-CLEVR reporters, Drosophila S2 cells were transfected with gypsy-CLEVR and PBS mutant constructs and imaged 48 hours post-transfection (Figure 1A-C). When co-transfected with tubulin-Gal4, required for the downstream expression of the WM markers, the gypsy-CLEVR reporter showed bright WM fluorescent signal in ∼3% of cells (Figure 1B and 1C). In contrast, no labeled cells were detected in the gypsy-CLEVR transfected cells when Gal4 was not present (Figure 1B and 1C). As previously reported (11), deletion or mutation of the primer binding site (gypsy-CLEVR^ΔPBS^, gypsy-CLEVR^PBSm2^) (Figure 1A) eliminated detection of WM labelled cells (Figure 1B and 1C). As controls to ensure a consistent rate of transfection, an actin5c-promoter driven WM dual reporter (pAc-WM), and a Gal4/UAS-driven WM plasmid (UAS-WM) (Figure 1A) were also transfected in parallel. pAc-WM, which does not require Gal4, displayed strong WM signal in ∼9% of cells, and UAS-WM labeled 0% and ∼10% of cells in the absence and presence of tubulin-Gal4 respectively (S1A and S1B Fig), consistent with our previously reported rates of S2 cells labeled with these constructs (11). Together, these findings confirm our previous report (Chang, Keegan, 2019) that gypsy-CLEVR labels S2 cells in which gypsy replication has occurred.

We next tested the gypsy-mCherry and gypsy-mCherry with *Env* deleted (gypsy-ΔEnv-mCherry) constructs to report gypsy expression when transfected into Drosophila S2 cells. Both of these constructs produce a nuclear localized mCherry signal when expressed (Figure 1B). We also tested the impact on gypsy-ΔEnv-mCherry when it was co-transfected with a pActin-driven gypsy-Env plasmid (pAc-Env) expressed in trans (Figure 1B). The gypsy-mCherry, gypsy-ΔEnv-mCherry, and gypsy-ΔEnv-mCherry co-transfected with pAc-Env each labeled ∼2% of cells (Figure 1C). For this set of experiments, we used a pActin-driven mCherry (pAc-H2B-mCherry) as a transfection control. The pAc-H2B-mCherry displayed a strong nuclear mCherry signal in ∼4% of cells (S1A and S1B Fig). Therefore, the gypsy-CLEVR and gypsy-mCherry groups of reporter constructs reliably label cells where gypsy has replicated or is expressed respectively, but these experiments do not discriminate between intercellular and intra-cellular replication cycles.

### CELL-ASSOCIATED TRANSMISSION OF GYPSY BETWEEN CO-CULTURED CELLS

We next used the gypsy-CLEVR reporter to test whether gypsy is capable of transmitting between cells grown in contact. We took advantage of the Gal4 dependence of the reporter expression in the gypsy-CLEVR construct. The gypsy-CLEVR reporter requires Gal4 to produce a fluorescent signal after replication, but does not require Gal4 for replication. We transfected separate populations of S2 cells with either tubulin-Gal4 or the gypsy-CLEVR reporter. 48 hours following transfection, cells were washed by centrifugation to remove remaining transfection complex, and then seeded into co-culture at equal ratios (Figure 2A). Cells were then mounted and imaged after 48 hours in co-culture. In this experiment, neither the Gal4 alone nor the gypsy-CLEVR alone is sufficient to yield expression of the dual WM reporter. On the other hand, intercellular transmission of the gypsy-CLEVR followed by integration into the Gal4 expressing recipient cell genome would yield reporter expression. As controls, we also used the gypsy-CLEVR^ΔPBS^ and gypsy-CLEVR^PBSm2^, which possess disrupted primer binding sites (Figure 2A) and therefore can be expressed but cannot replicate. We also used a co-culture control in which one population of cells had been transfected with a Gal4 dependent UAS-WM and the other with the Gal4 itself. The expectation is that there should be no intercellular transmission of the WM transcript when it is not associated with the gypsy-CLEVR construct.

**Figure 2:**
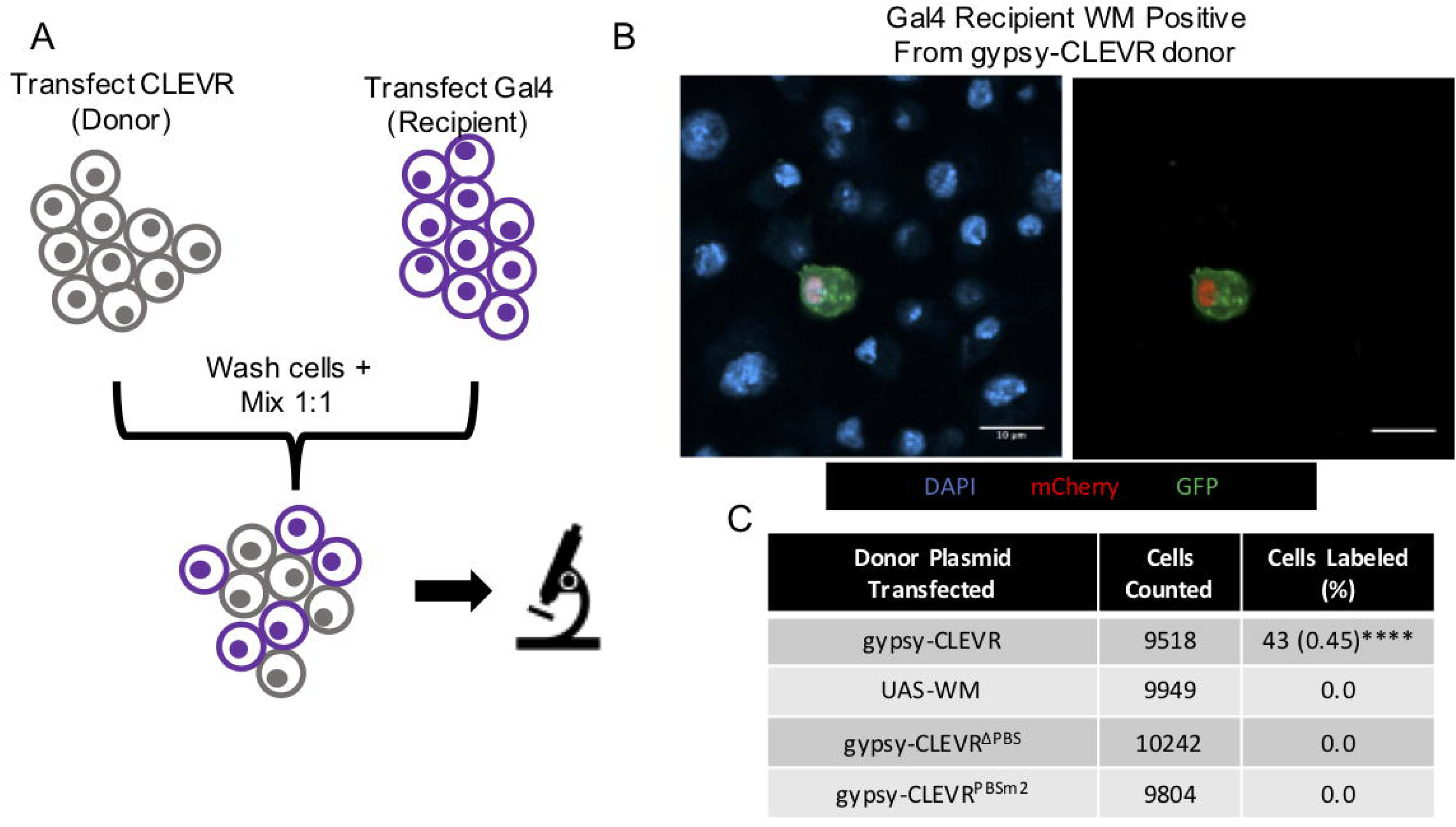
The gypsy-CLEVR reporter reveals that gypsy transfers between cells in contact and integrates into the infected recipient cell. (A) Cartoon schematic showing the experimental design of the co-culture assay. Separate populations of S2 cells are transfected with gypsy-CLEVR or tubulin-Gal4 constructs, washed, and then mixed together in equal proportions. (B) Fluorescent images showing WM labeled cells in the co-cultured gypsy-CLEVR and tubulin-Gal4 cell population. UAS-WM, gypsy-CLEVR^ΔPBS^, and gypsy-CLEVR^PBSm2^ showed no WM labeled cells and are not shown. Scale bars = 10 µm. (C) Quantification showing the percentage of cells expressing the WM reporter for the UAS-WM (control) and gypsy-CLEVR constructs in co-culture with tubulin-Gal4. Quantification is presented as totals cells counted from 3 near equivalent sets of biological replicates. Significance was calculated against UAS-WM. Significance was determined using the Fisher’s Exact test variant of the Chi^2^ test. Significance values are denoted as: p=<0.05 *, p=<0.001 ***, p=<0.0001****

We see clear evidence that gypsy is able to transmit between cells in this cell-associated co-culture assay. When the intact gypsy-CLEVR construct was used, it resulted in positive WM expression detected in ∼0.5% of cells, indicating gypsy containing the properly rearranged UAS-WM reporter CLEVR system is capable of moving into tubulin-Gal4 expressing cells (Figure 2B and 2C). In contrast, we observed no WM positive cells when the UAS-WM transfected cells were co-cultured with Gal4 transfected cells, indicating that the reporter cannot move between cells when it is not associated with gypsy. In addition, we observe no WM positive label when gypsy-CLEVR^ΔPBS^ or gypsy-CLEVR^PBSm2^ transfected cells were co-cultured with Gal4 expressing cells (Figure 2C). Thus, gypsy constructs that are unable to replicate due to deletion or mutation of the PBS also are unable to transmit between co-cultured cells grown in contact.

### CELL-FREE TRANSMISSION OF GYPSY

We next tested whether gypsy is capable of cell-free transmission between S2 cells that are not grown in direct cell contact. This assay is conceptually similar to that of the gypsy-CLEVR reporter in co-culture described above. However, in this case, we used a transwell system that utilizes a semi-permeable barrier (0.4 µm) between two separately transfected populations of cells. In a manner similar to that of the co-culture assay, we capitalized on the Gal4 dependence of the WM reporter in the gypsy-CLEVR construct. This construct is capable of replicating independently of Gal4, but cannot express the reporter from the integrated pro-virus unless Gal4 is present. We again separately transfected either the gypsy-CLEVR reporter or Gal4, and we grew these in a transwell cell culture plate to separate the two populations of cells. The culture plates used possess a membrane permeabilized by 0.4 µm pores, which are sufficient to restrict passage of whole cells, the nuclei of which are several microns in diameter, and likely most cellular debris, but would permit transfer of virus particles that likely would be below that size.

Here too, we tested transmission of the wild-type gypsy-CLEVR as well as the gypsy-CLEVR^ΔPBS^, and gypsy-CLEVR^PBSm2^, which are unable to replicate due to disruption of the PBS sequences. A separate population of cells was transfected with Gal4 alone. Cells transfected with either the CLEVR constructs or the Gal4 were allowed to incubate on their own for 48 hours, after which the cells were washed by centrifugation and seeded on opposite sides of the membrane in the transwell plate cell-culture dish (Figure 3A). The gypsy-CLEVR transfected populations were designated as “donor cells” while the tubulin-Gal4 transfected populations were designated as “recipient cells”. After an additional incubation of 48 hours in the transwell cell culture plate, both donor and recipient populations were separately mounted and imaged to detect both transfer and directionality of transfer. Expression of the WM reporter was indicative of transfer, as none of the plasmids transfected can produce the WM signal on their own.

**Figure 3:**
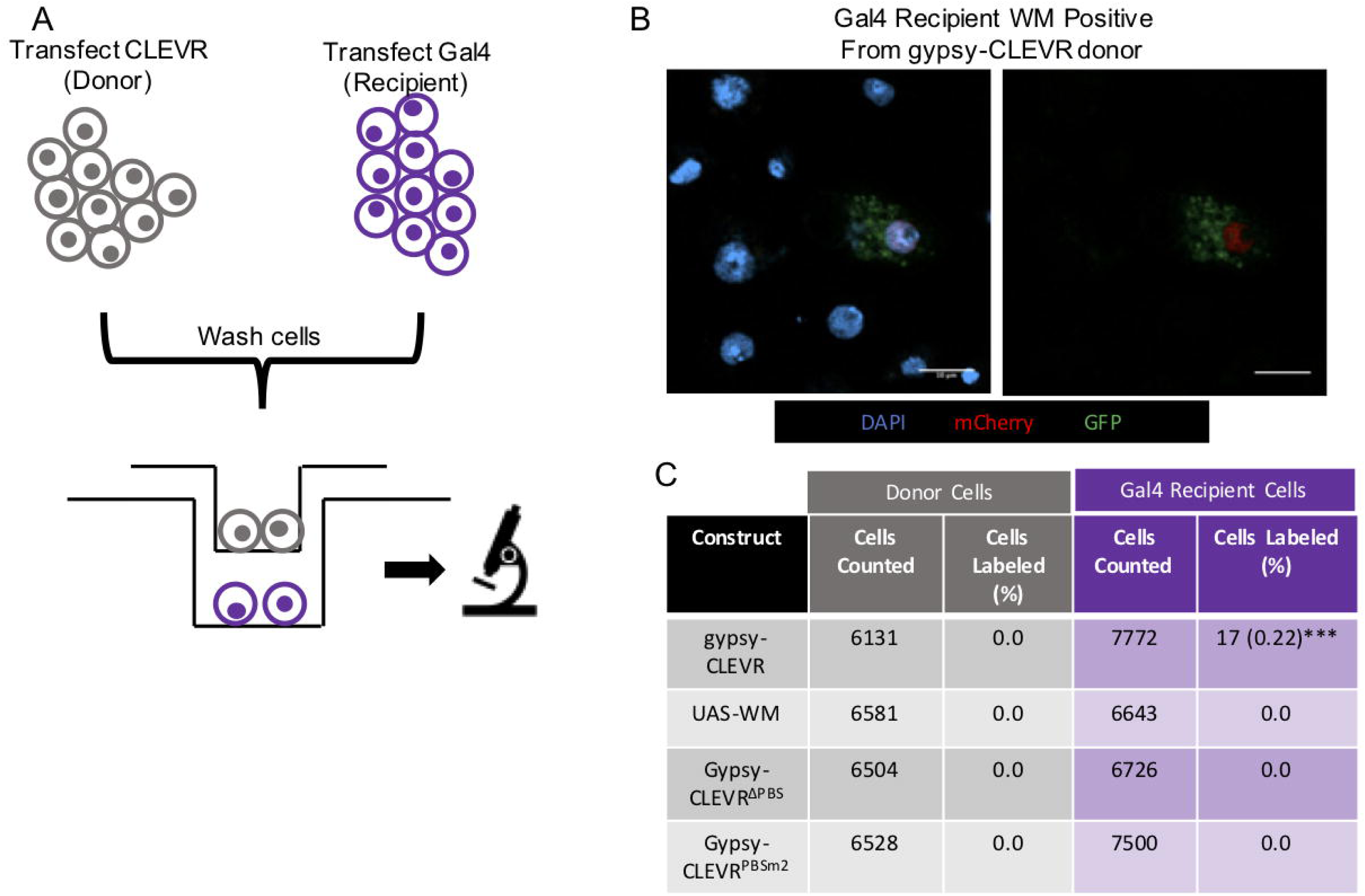
The gypsy-CLEVR reporter reveals intercellular gypsy transmission through a contact restricting membrane. (A) Cartoon schematic showing the experimental design of the transwell assay. Separate populations of S2 cells are transfected with gypsy-CLEVR or tubulin-Gal4 constructs, washed, and then re-seeded on separate sides of a 0.4 µm membrane. (B) Fluorescent images showing WM labeled cells in the tubulin-Gal4 cell recipient population. UAS-WM, gypsy-CLEVR^ΔPBS^, and gypsy-CLEVR^PBSm2^ showed no WM labeled cells in the recipient populations and are not shown. No WM labeled cells were detected in the donor populations and are not shown. Scale bars = 10 µm. (C) Quantification showing the percentage of cells expressing the WM reporter for the UAS-WM (control) and gypsy-CLEVR constructs for both the donor and recipient populations in the transwell assay. Quantification is presented as totals cells counted from 3 near equivalent sets of biological replicates. Significance was calculated against UAS-WM. Significance was determined using the Fisher’s Exact test variant of the Chi^2^ test. Significance values are denoted as: p=<0.05 *, p=<0.001 ***, p=<0.0001****

Among all the groups, the only population of cells that displayed WM dual fluorescence signal were the “recipient” population of cells expressing tubulin-Gal4 when they were grown on the opposite side of the membrane to the intact gypsy-CLEVR donor population (Figure 3). In this recipient population of cells, ∼0.2% of Gal4 transfected cells were found to express the WM reporter (Figure 3B and 3C). No donor populations (gypsy-CLEVR, gypsy-CLEVR^ΔPBS^, gypsy-CLEVR^PBSm2^) or the donor control (UAS-WM) displayed any WM signal, indicating that Gal4 was in no case transferred across the membrane from the recipient to the donor cells (Figure 3C). Further, we did not observe any WM reporter expression in the tubulin-Gal4 recipient populations grown opposite the gypsy-CLEVR^ΔPBS^, gypsy-CLEVR^PBSm2^, that are unable to replicate (Figure 3C). Nor did we observe any expression in the Gal4-expressing recipient cells grown in the transwell below the UAS-WM control donor populations (Figure 3C). Together, these findings demonstrate that gypsy is able to transmit between cells that are not in contact. The fact that such transfer only occurs when the reporter is tethered to an intact gypsy that is able to replicate demonstrates the specificity of this assay. The unidirectional nature of transfer from gypsy-CLEVR expressing cells to tubulin-Gal4 expressing recipient cells, also supports the conclusion that gypsy acts as an infectious retrovirus in cell-culture, capable of cell-free transmission.

### INTERCELLULAR TRANSMISSION OF GYPSY REQUIRES ENV

Enveloped viruses encode a surface glycoprotein that mediates recognition of cellular receptors and fusion with the cell membrane. Retroviral *Env* genes thus are required both for cell-free and cell-associated transmission. To test whether intercellular transmission of gypsy also is env-dependent, we used the gypsy-ΔEnv-mCherry construct, in which we replaced the gypsy-encoded env ORF with that of mCherry. We tested both the gypsy-mCherry with *Env* intact (Figure 1A) and gypsy-ΔEnv-mCherry constructs in the transwell assay that is described above for the gypsy-CLEVR reporter. Unlike the WM reporter in gypsy-CLEVR assay, the expression of mCherry from gypsy-mCherry and gypsy-ΔEnv-mCherry does not require replication of the gypsy RNA genome and does not require co-expression of Gal4.

S2 cells were transfected with either gypsy-mCherry or gypsy-ΔEnv-mCherry. As a further test of the requirement for Env, we also tested whether co-transfection of a pActin-Env (pAc-Env) was able to rescue the env-deficient virus in trans. A pAc-H2B-mCherry plasmid was used as a transfection control. As with the gypsy-CLEVR system described above, transfected cells were first cultured separately for 48 hours (Figure 4A). Following this incubation period, the cells were washed via centrifugation and transferred into the transwell cell culture plate above recipient S2 cells (Figure 4A). Because the gypsy-mCherry constructs do not require presence of Gal4 to visualize reporter expression, the recipient cells used here were untransfected. This offers a numerical advantage over the gypsy-CLEVR reporter in that 100% of the recipient pool of cells are able to report transmission if it occurs. Following a 48-hour incubation in the transwell cell culture plate, both donor and recipient populations of cells were mounted and imaged for expression of nuclear mCherry.

**Figure 4:**
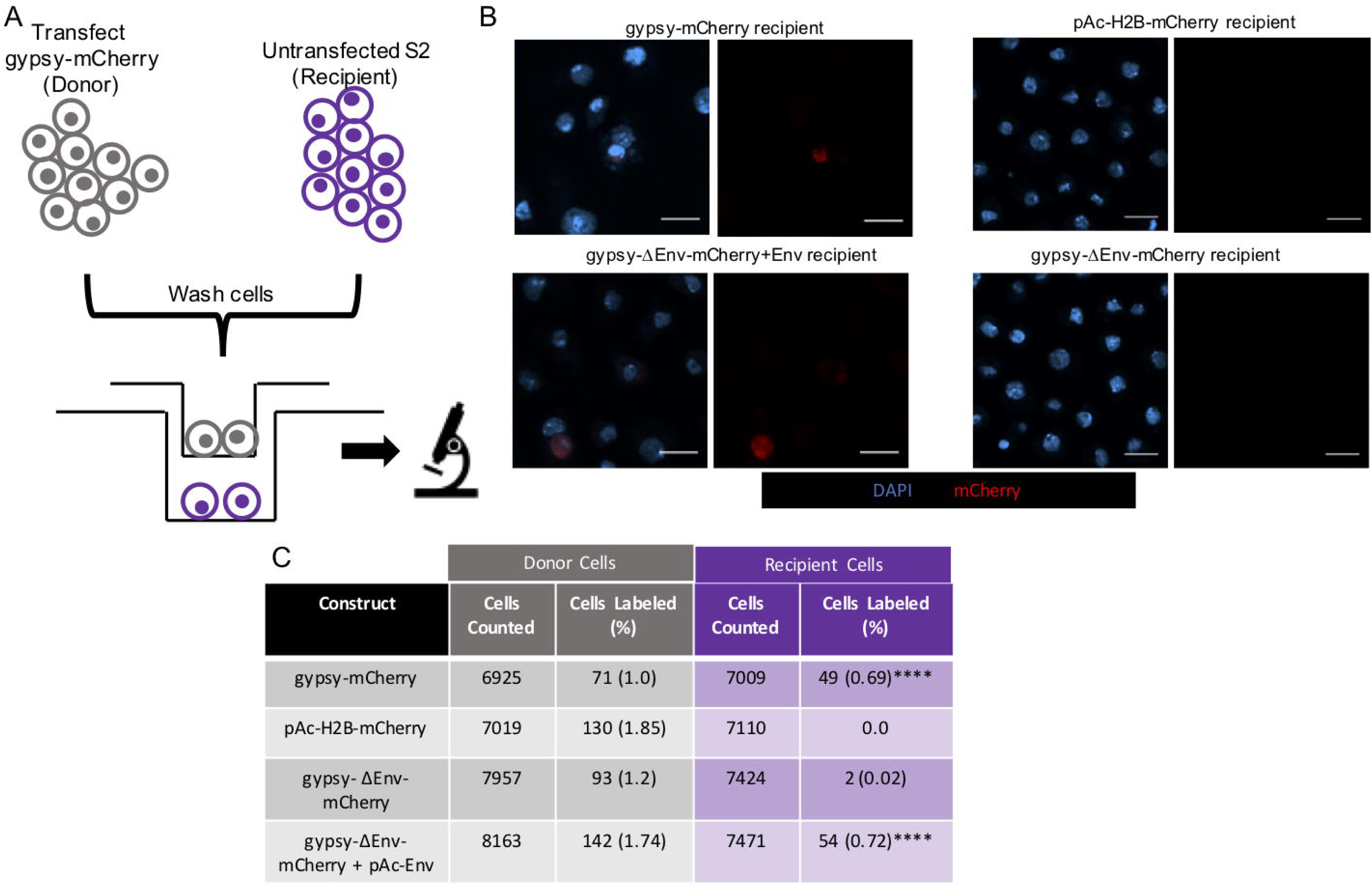
The gypsy-mCherry reporter reveals that intercellular transmission of gypsy requires functional env. A) Cartoon schematic showing the experimental design of the transwell assay. One population is transfected with the gypsy-mCherry constructs, washed, and placed opposite untransfected S2 cells separated by a 0.4 µm membrane. (B) Fluorescent images showing mCherry labeled cells in the S2 cell recipient population. pAc-H2B-mCherry and gypsy-ΔEnv-mCherry recipient populations show no or few labeled cells respectively, and are not shown. Donor populations are not shown. Scale bars = 10 µm. (C) Quantification showing the percentage of cells expressing mCherry for the pAc-H2B-mCherry (control) and gypsy-mCherry constructs for both the donor and recipient populations in the transwell assay. Quantification is presented as totals cells counted from 3 near equivalent sets of biological replicates. Significance was calculated against gypsy-ΔEnv-mCherry. Significance was determined using the Fisher’s Exact test variant of the Chi^2^ test. Significance values are denoted as: p=<0.05 *, p=<0.001 ***, p=<0.0001****

Here, all of the transfected donor population are expected to express nuclear mCherry, and the recipient population of cells would express mCherry if gypsy had transferred across the membrane. Within the donor populations of cells, the control pAc-H2B-mCherry plasmid showed expression that labeled ∼2% of cells, reflecting the transfection rate at this time-point (4 days after transfection). The percent of mCherry-expressing donor cells for gypsy-mCherry, gypsy-ΔEnv-mCherry, and gypsy-ΔEnv-mCherry + pAc-Env transfections were ∼1%, ∼1% and ∼2% respectively (Figure 4B and 4C). In the recipient population grown opposite to the control pAc-H2B-mCherry, no cells were found to express the mCherry label, as expected. In contrast, ∼0.7% of recipient cells grown opposite to the gypsy-mCherry were found to express the reporter, consistent with the fact that gypsy virus can transmit between cells that are not in contact. But this number dropped to near zero (0.02%) for recipient cells grown opposite to gypsy-ΔEnv-mCherry transfected donor cells. This strongly supports the conclusion that the gypsy *Env* gene is required for transmission. This deficiency in intercellular transmission with the *Env* deleted construct also could be rescued when Env was expressed in trans. When the gypsy-ΔEnv-mCherry construct was co-transfected with pAc-Env, ∼0.7% of the recipient cells expressed mCherry (Figure 4B and 4C). Together, these results confirm that gypsy is capable of cell-free transmission, but also show that this transmission is reliant upon the presence of functional *Env*.

### INTRACELLULAR TRANSPOSITION OF GYPSY IS ENV INDEPENDENT

The above findings indicate that gypsy retains the ability to transmit between cells under both cell-associated and cell-free conditions, and such transmission is *Env* dependent. Unlike retroviruses, LTR-retrotransposons typically utilize an intracellular replication cycle that is not env-dependent, but intracellular replication also requires significant differences in targeting within the cell. The *Env* dependent inter-cellular transmission of gypsy would necessitate assembly at the extracellular membrane. We wondered therefore if gypsy, which classically has been thought of as a retrotransposon, is even capable of replicating intracellularly. To test this, we generated a gypsy-CLEVR construct in which we had introduced a frameshift mutation within the Env ORF. Because the gypsy-CLEVR reporter labels cells only after reverse transcription and template switching (10, 11), this reporter provides a means to distinguish replication events from mere expression. Because expression of the WM dual reporter that is contained on the gypsy-CLEVR construct is Gal4 dependent, we co-transfected with Gal4 expression construct.

S2 cells were transfected with Tubulin-Gal4 as well as either gypsy-CLEVR or gypsy-CLEVR^ΔEnv^. Each of the above two constructs were tested both with and without pAc-Env to provide Env expression in trans (Figure 5A). After transfection, the populations of cells were incubated for 48 hours, mounted and imaged to detect the presence of the WM reporter. When co-transfected with tubulin-Gal4, the gypsy-CLEVR plasmid produced strong WM label in 3.2% of cells imaged. This is consistent with the robust levels of gypsy replication in S2 cells that we have observed previously (11). When this gypsy-CLEVR construct was co transfected with both tubulin-Gal4 and an additional source of pAc-Env expressed in trans, the fraction of labeled cells remained at 3.2% (Figure 5B and 5C). Thus, Env levels are not limiting the rate of replication of the gypsy-CLEVR construct. When gypsy-CLEVR^ΔEnv^ was co-transfected with tubulin-Gal4, 3.0% of cells were WM labeled, a rate that is statistically indistinguishable from that of the intact gypsy-CLEVR construct. Similarly, when the gypsy-CLEVR^ΔEnv^ was co-transfected with both tubulin-Gal4 and pAc-Env, 3.1% of cells were labeled with WM (Figure 5B and 5C). Here too, the rate of replication of gypsy-CLEVR^ΔEnv^ is indistinguishable from that of the intact gypsy-CLEVR irrespective of whether an additional source of Env is supplied in trans. Taken together, these findings demonstrate that gypsy is capable of intracellular retrotransposition that is independent of Env.

**Figure 5:**
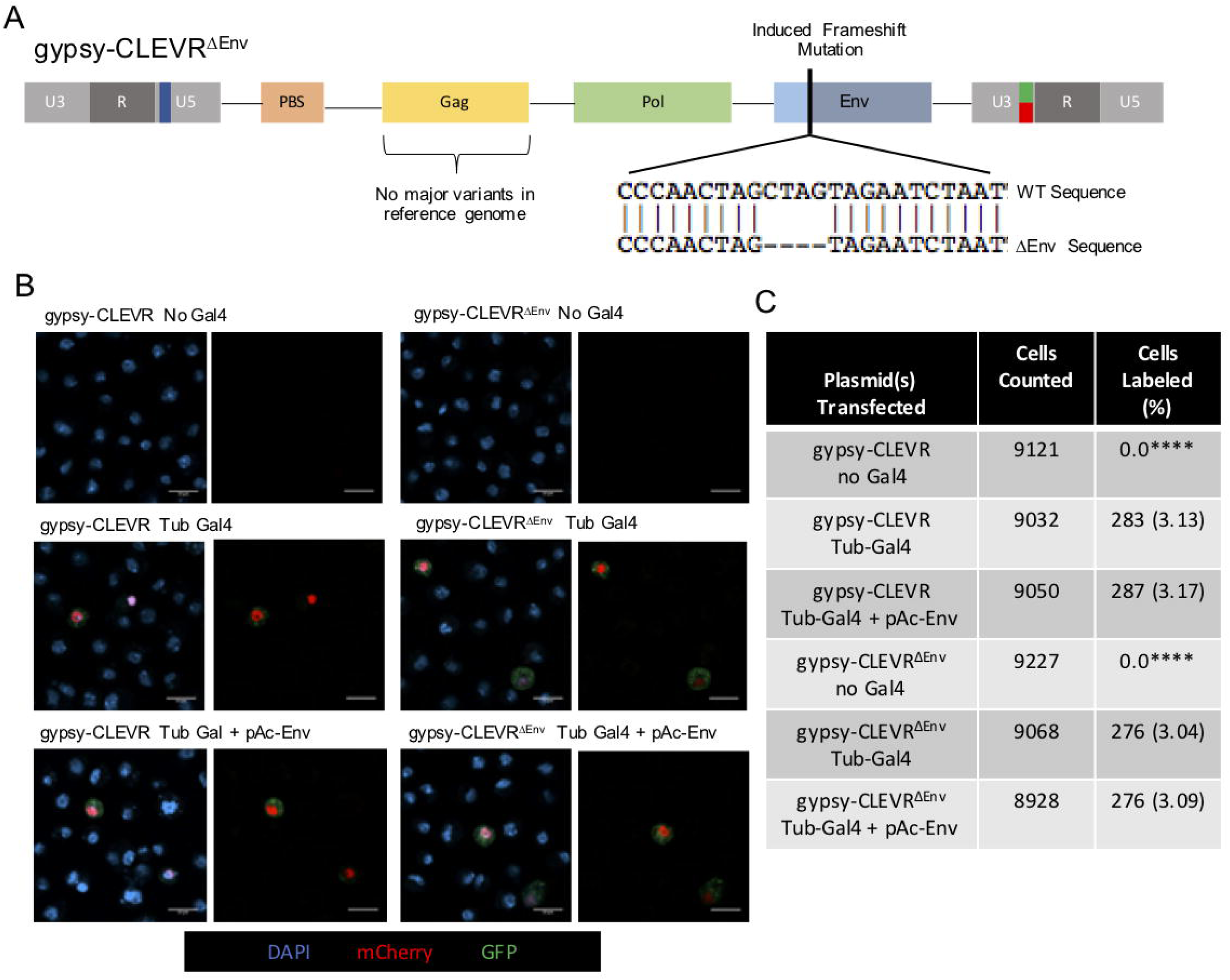
Functional env is not required for the intracellular retrotransposition of gypsy. A) Cartoon schematic showing the overall structure of the gypsy-CLEVR^ΔEnv^ construct, which is identical to gypsy-CLEVR but has a frameshift mutation within the env ORF, seen in detail in the sequence comparison directly below. B) Fluorescent images showing the absence of WM signal in gypsy-CLEVR and gypsy-CLEVR^ΔEnv^ populations lacking Gal4, and positive WM signal in gypsy-CLEVR and gypsy-CLEVR^ΔEnv^ when co-transfected with tubulin Gal4 as well as with pAc-Env. C) Quantification of the percentage of cells that showed positive WM signal for gypsy-CLEVR and gypsy-CLEVR^ΔEnv^ with and without Gal4, as well as with pAc-Env. No statistically significant differences were found absent gypsy-CLEVR and gypsy-CLEVR^ΔEnv^ lacking the presence of Gal4. Quantification is presented as totals cells counted from 3 near equivalent sets of biological replicates. Significance was determined using the Fisher’s Exact test variant of the Chi^2^ test. Significance values are denoted as: p=<0.05 *, p=<0.001 ***, p=<0.0001****

## Discussion

ERVs can defy a clear definition, as some act as retroviruses and others act as LTR-RTEs, leaving these elements in a sort of evolutionary “gray area”. From an evolutionary perspective, it is thought that LTR-RTEs are the likely ancestors of retroviruses, and all vertebrate retroviruses come from a single lineage (85, 99). The emergence of retroviruses is thought to have involved a multi-step process that includes targeting to the cell membrane and the incorporation of a surface glycoprotein (Env). This process also has likely occurred in reverse, as some ERVs have lost their Env and developed the ability to re-target internally, and in some cases, these have even been called RTEs, despite the different evolutionary history. Indeed, the fact that the invertebrate gypsy element contains an *env* gene is suggestive that the gypsy element is capturing the process of the generation of a new lineage of retrovirus from RTE ancestors.

Here, we demonstrate that gypsy, one of the most well-known LTR-RTEs, is able to replicate intracellularly as an RTE, but also can transmit between cells grown in culture as a virus. This intercellular transmission can occur both for cells grown in close contact and by a cell free mechanism. We observe such intercellular movement with two different reporters, one of which labels any recipient cells that express gypsy encoded proteins and the other of which only labels recipient cells that have a gypsy provirus which has gone through reverse transcription. With this second reporter, we only observe transmission when the PBS, which is essential for viral replication, is intact. Finally, we demonstrate that intercellular transmission of gypsy occurs by a mechanism that requires functional *Env*, consistent with the idea that transmission occurs via a viral mechanism. It is worth noting that it has previously been suggested that a tagged gypsy element may be capable of transmission in vivo from somatic follicle cells to the oocyte, and that this may occur in the absence of a functional Env (100). Although we do not observe *Env* independent intercellular transmission in S2 cells, we cannot rule out the possibility that in some contexts, gypsy exhibits a third mode of replication that is *Env* independent but intercellular.

The gypsy element, which has been termed an errantivirus, has long been thought to possess features of an infectious retrovirus (92, 101). Pseudotyping of Moloney murine leukemia virus with gypsy env is sufficient to confer entry insect cells (102), which demonstrates that the gypsy envelope glycoprotein is functional. Several reports document that virus-like particles are present in Drosophila ovaries from genotypes in which gypsy replication is taking place (93, 94). More striking is the observation that horizontal transmission of gypsy can occur when larvae from strains that have no functional gypsy elements are fed extracts from ovaries of animals with active gypsy (91, 94). The experiments that we describe here demonstrate that gypsy indeed possesses qualities of a retrovirus, enabling Env-dependent infectious transmission. More surprisingly, gypsy also can replicate just as efficiently as an intracellular RTE in the absence of Env. Given the complex functional changes that underlie evolutionary transition between LTR-RTEs and retroviruses, this dual mode of replication is unexpected. This point is driven home by a comparison to the murine intracisternal A-type particle (IAP) and the related intracisternal A-type particle with Env (IAP-E).

The IAP elements, which are murine ERVs that lack env, follow a purely intracellular RTE-like replication lifecycle, remaining within the cell where they are targeted to the lumen of the endoplasmic reticulum, which is contiguous with the perinuclear space (81). Conversely, the mouse IAP-E element, which possesses a functional env ORF and is therefore more closely related to the ancestral exogenous virus that gave rise to all IAP and IAP-E ERVs, has been shown to replicate following an intercellular lifecycle, producing exogenous virus that buds at the membrane and infects neighboring cells (81). Although the loss of *Env* is important in the evolutionary transition from the viral life cycle into to an RTE-like lifecycle, mouse IAP and IAP-E elements also differ in the gag ORF, where amino acid variation within the gag proteins of these elements are sufficient to change the targeting to be compatible either with intercellular or intracellular replication (81). Strikingly, hybrid IAP-E elements in which the N-terminal region of gag is substituted from IAP are unable to produce viral particles at the membrane because of mis-targeting of gag.

The situation with gypsy appears to be quite different from that of IAP-E. Unlike these murine elements, all of the intact gypsy copies that are identified in the *Drosophila* reference genome appear to contain an *Env* reading frame. And we see no evidence for existence of gypsy variants with significant substitutions in gag that might provide for two classes of element as is the case with IAP/IAP-E. Moreover, in contrast with IAP/IAP-E, the specific variant of gypsy that we used to construct our reporters appears capable of both modes of replication. The ability of ERVs to replicate via intracellular vs intercellular mechanisms may have significant biological impact.

Expression and replication of RTEs and ERVs have been found in somatic tissues both during normal development (5, 7, 8, 11-13, 16, 17, 19, 21-24, 83, 103, 104), in advanced aging (11, 16, 27, 28, 33, 34, 86, 105-107) and in diseases of aging such as neurodegeneration (10, 57-61, 65, 67, 72, 96, 108-110), and cancer (9, 48-52, 111-118). The functional consequences of somatic expression and replication of RTEs/ERVs are only beginning to be understood, and it is not known if inter-cellular transmission occurs in vivo. But there already is evidence that cells that exhibit RTE/ERV replication may have non-cell autonomous impacts on surrounding tissue (10, 28). It now is established that HERV-K (96, 97), IAP-E (81, 95) and gypsy each are functional viruses in cell culture and IAP (81) and gypsy have intracellular replication cycles as well. While LINE elements do not encode machinery for viral transmission, there is recent evidence that human-specific LINE-1 elements can transmit between cells in culture via extracellular vesicles (79). In addition, the Arc genes in both mammals and in *Drosophila* have recently been found to have their ancestral origin from a gypsy-family gag protein, and Arc been shown to bind and transport mRNA cargo between neurons (78, 80). Together, these findings reveal the dual replication strategies used by an element in transition between a retrotransposon and a virus and raise the possibility that ERVs and RTEs may provide routes for transfer of genetic information between cells within an organism.

## Materials and Methods

### Constructs

To generate pAc-Env, the Env was amplified from the gypsy-CLEVR plasmid using polymerase chain reaction (PCR) and was inserted into the multiple cloning site (MCS) of the pAc5.1 C vector (Thermo Fisher Scientific) with a NotI and KpnI digestion. The gypsy-CLEVR^ΔEnv^ was constructed by digesting the gypsy-CLEVR plasmid with BcuI, ethanol precipitated, and treated with Klenow before ligation, resulting in a frame shift occurring within the Env of gypsy-CLEVR at position 13,847 in the CLEVR reporter. Gypsy Env is located between 13,470-14,916 within the CLEVR construct. To generate the S2 cell-based reporter pAc-H2B-mCherry, the nuclear localization reporter H2B-mCherry-HA was amplified from Watermelon (WM) reporter described in our previous study (11) by PCR. The PCR-amplified H2B-mCherry-HA was then inserted into the XhoI site of the *Drosophila* constitutive expression vector, pAc5.1/V5-His version C (V411020, Thermo Fisher Scientific). In order to test the transferring ability of gypsy, the gypsy backbone used in previous publication (11) was amplified and cloned into the NotI/XbaI sites of pAc5.1/V5-His version B (V411020, Thermo Fisher Scientific). To synthesize the final pAc-gypsy-H2B-mCherry vector, the H2B-mCherry-HA DNA fragment from WM was then added to the end of gypsy ORF3 (env) by P2A linking peptide sequences, with the stop codon of gypsy ORF3 (env) removed. To synthesize pAc-gypsy-H2B-mCherry^ΔEnv^, the whole gypsy ORF3 (env) was deleted from gypsy backbone and replaced with H2B-mCherry-HA fragment, except these initial <u>AG</u>**G**TTCACCCTCATG nucleotides from env were maintained in order to provide the endogenous splicing accepting site to receive the alternative splicing stat codon **AT**<u>GT</u> from gypsy ORF1 (gag) (93).

### Cell Culture

Drosophila S2 cells (R69007, Thermo Fisher Scientific) were cultured in *Schneider’s Drosophila Media* (Thermo Fisher Scientific) supplemented with 10% Fetal Bovine Serum (Thermo Fisher Scientific) and Pennicillin-Streptomycin-Glutamine (Thermo Fisher Scientific), in 75cm^2^ flasks (Flask info). Cells were transfected with 1.5ug of each plasmid DNA with the Effectene transfection kit (Qiagen). After 48 hours in transfection complex, cells were fixed in 4% Paraformaldehyde and mounted on coverslips coated in 0.5mg/ml Concanavalin A and ProLong Diamond Antifade Mountant with DAPI (Thermo Fisher Scientific). All images were taken on a Zeiss Confocal microscope and quantified under blinded conditions using the cell counter feature in FIJI.

### Co-Culture

Prior to co-culture, cells were transfected with individual plasmids and incubated for 48 hours in transfection complex. Following this incubation, cells were washed 3 times with 5ml of Schneider’s Drosophila Media and co-cultured at a 50:50 ratio in 75cm^2^ flasks. After 48 hours in the co-culture condition, cells were fixed in 4% Paraformaldehyde and mounted on coverslips coated in 0.5mg/ml Concanavalin A and ProLong Diamond Antifade Mountant with DAPI.

### Transwell

Prior to introduction into the Transwell system, cells were transfected with individual plasmids and incubated for 48 hours in transfection complex. Following this incubation, cells were washed 3 times with 5ml of Schneider’s *Drosophila* Media and recipient and donor cells were moved to opposing sides of a 6-well, 0.4um Transwell plate. Following 48 hours in the Transwell plate, cells from both sides of the plate were individually fixed in 4% Paraformaldehyde and mounted on coverslips coated in 0.5mg/ml Concanavalin A and ProLong Diamond Antifade Mountant with DAPI.

### Statistical Analysis

All data was analyzed using the Chi^2^ with Yate’s correction analysis in order to obtain a P value for significance between separate groups. For comparisons incorporating multiple zeroes, the Fisher’s Exact test variant of the Chi^2^ test was used. Significance values are denoted as: p=<0.05 *, p=<0.01**, p=<0.001 ***, p=<0.0001****

### PCR primers used

The following primers were used to amplify gypsy Env from the gypsy-CLEVR construct:

F: GGTACCCAAAACATGatGTTCACCCTCATGATGTTCATACC

R:GGGAGTAGTTAACAACTAAGCGGCCGCAATTTAGCGCGC

Reverse complement of the R primer:

GCGCGCTAAATTGCGGCCGCTTAGTTGTTAACTACTCCC

## Acknowledgements

We are thankful to all of the members of the Dubnau and Metzger research groups for helpful discussions and comments on the manuscript. This work was supported by an NIH K22 CA226047 award to M. Metzger and NINDS award R01NS091748 and NIA award RF1AG057338 to J. Dubnau.

---

**Supplemental Figure 1: Control plasmids function in Drosophila S2 cell culture**

A) Fluorescent images showing mCherry labeled nuclei present in approximately 4% of pAc-H2B-mCherry transfected cells, as well as WM signal expressed in approximately 9.6% and 10.3% of pAc-WM and UAS-WM cotransfected with Tub Gal4 transfected cells respectively. UAS-WM, when not cotransfected with a Gal4 plasmid showed no expression of the WM reporter.

B) Quantification of the cells counted and the percentage and number (in parentheses) of cells expressing the control pAc-H2B-mCherry, pAc-WM, and UAS-WM without and without Gal4 plasmids. Quantification is presented as totals cells counted from 3 near equivalent sets of biological replicates.

## References

1. Huang CR, Burns KH, Boeke JD. Active transposition in genomes. Annu Rev Genet. 2012;46:651–75.

2. Kazazian HH, Jr. Mobile elements: drivers of genome evolution. Science. 2004;303(5664):1626–32.

3. Babushok DV, Kazazian HH, Jr. Progress in understanding the biology of the human mutagen LINE-1. Hum Mutat. 2007;28(6):527–39.

4. Dubnau J. The Retrotransposon storm and the dangers of a Collyer’s genome. Curr Opin Genet Dev. 2018;49:95–105.

5. Faulkner GJ, Billon V. L1 retrotransposition in the soma: a field jumping ahead. Mob DNA. 2018;9:22.

6. Reilly MT, Faulkner GJ, Dubnau J, Ponomarev I, Gage FH. The role of transposable elements in health and diseases of the central nervous system. J Neurosci. 2013;33(45):17577–86.

7. Richardson SR, Morell S, Faulkner GJ. L1 retrotransposons and somatic mosaicism in the brain. Annu Rev Genet. 2014;48:1–27.

8. Baillie JK, Barnett MW, Upton KR, Gerhardt DJ, Richmond TA, De Sapio F, et al. Somatic retrotransposition alters the genetic landscape of the human brain. Nature. 2011;479(7374):534–7.

9. Burns KH. Transposable elements in cancer. Nat Rev Cancer. 2017;17(7):415–24.

10. Chang YH, Dubnau J. The Gypsy Endogenous Retrovirus Drives Non-Cell-Autonomous Propagation in a Drosophila TDP-43 Model of Neurodegeneration. Curr Biol. 2019;29(19):3135–52 e4.

11. Chang YH, Keegan RM, Prazak L, Dubnau J. Cellular labeling of endogenous retrovirus replication (CLEVR) reveals de novo insertions of the gypsy retrotransposable element in cell culture and in both neurons and glial cells of aging fruit flies. PLoS Biol. 2019;17(5):e3000278.

12. Coufal NG, Garcia-Perez JL, Peng GE, Yeo GW, Mu Y, Lovci MT, et al. L1 retrotransposition in human neural progenitor cells. Nature. 2009;460(7259):1127–31.

13. Eickbush MT, Eickbush TH. Retrotransposition of R2 elements in somatic nuclei during the early development of Drosophila. Mob DNA. 2011;2(1):11.

14. Kazazian HH, Jr., Moran JV. Mobile DNA in Health and Disease. N Engl J Med. 2017;377(4):361–70.

15. Kubo S, Seleme MC, Soifer HS, Perez JL, Moran JV, Kazazian HH, Jr., et al. L1 retrotransposition in nondividing and primary human somatic cells. Proc Natl Acad Sci U S A. 2006;103(21):8036–41.

16. Li W, Prazak L, Chatterjee N, Gruninger S, Krug L, Theodorou D, et al. Activation of transposable elements during aging and neuronal decline in Drosophila. Nat Neurosci. 2013;16(5):529–31.

17. Muotri AR, Chu VT, Marchetto MC, Deng W, Moran JV, Gage FH. Somatic mosaicism in neuronal precursor cells mediated by L1 retrotransposition. Nature. 2005;435(7044):903–10.

18. Bae BI, Jayaraman D, Walsh CA. Genetic changes shaping the human brain. Dev Cell. 2015;32(4):423–34.

19. Evrony GD, Cai X, Lee E, Hills LB, Elhosary PC, Lehmann HS, et al. Single-neuron sequencing analysis of L1 retrotransposition and somatic mutation in the human brain. Cell. 2012;151(3):483–96.

20. Poduri A, Evrony GD, Cai X, Walsh CA. Somatic mutation, genomic variation, and neurological disease. Science. 2013;341(6141):1237758.

21. Cai X, Evrony GD, Lehmann HS, Elhosary PC, Mehta BK, Poduri A, et al. Single-cell, genome-wide sequencing identifies clonal somatic copy-number variation in the human brain. Cell Rep. 2014;8(5):1280–9.

22. Bedrosian TA, Linker S, Gage FH. Environment-driven somatic mosaicism in brain disorders. Genome Med. 2016;8(1):58.

23. Muotri AR, Marchetto MC, Coufal NG, Oefner R, Yeo G, Nakashima K, et al. L1 retrotransposition in neurons is modulated by MeCP2. Nature. 2010;468(7322):443–6.

24. Muotri AR, Zhao C, Marchetto MC, Gage FH. Environmental influence on L1 retrotransposons in the adult hippocampus. Hippocampus. 2009;19(10):1002–7.

25. Faulkner GJ. Retrotransposons: mobile and mutagenic from conception to death. FEBS Lett. 2011;585(11):1589–94.

26. De Cecco M, Criscione SW, Peckham EJ, Hillenmeyer S, Hamm EA, Manivannan J, et al. Genomes of replicatively senescent cells undergo global epigenetic changes leading to gene silencing and activation of transposable elements. Aging Cell. 2013;12(2):247–56.

27. De Cecco M, Criscione SW, Peterson AL, Neretti N, Sedivy JM, Kreiling JA. Transposable elements become active and mobile in the genomes of aging mammalian somatic tissues. Aging (Albany NY). 2013;5(12):867–83.

28. De Cecco M, Ito T, Petrashen AP, Elias AE, Skvir NJ, Criscione SW, et al. L1 drives IFN in senescent cells and promotes age-associated inflammation. Nature. 2019;566(7742):73–8.

29. Driver CJ, McKechnie SW. Transposable elements as a factor in the aging of Drosophila melanogaster. Ann N Y Acad Sci. 1992;673:83–91.

30. Elsner D, Meusemann K, Korb J. Longevity and transposon defense, the case of termite reproductives. Proc Natl Acad Sci U S A. 2018;115(21):5504–9.

31. St Laurent G, 3rd, Hammell N, McCaffrey TA. A LINE-1 component to human aging: do LINE elements exact a longevity cost for evolutionary advantage? Mech Ageing Dev. 2010;131(5):299–305.

32. Wood JG, Helfand SL. Chromatin structure and transposable elements in organismal aging. Front Genet. 2013;4:274.

33. Wood JG, Jones BC, Jiang N, Chang C, Hosier S, Wickremesinghe P, et al. Chromatin-modifying genetic interventions suppress age-associated transposable element activation and extend life span in Drosophila. Proc Natl Acad Sci U S A. 2016;113(40):11277–82.

34. Maxwell PH, Burhans WC, Curcio MJ. Retrotransposition is associated with genome instability during chronological aging. Proc Natl Acad Sci U S A. 2011;108(51):20376–81.

35. Maxwell PH, Curcio MJ. Incorporation of Y’-Ty1 cDNA destabilizes telomeres in Saccharomyces cerevisiae telomerase-negative mutants. Genetics. 2008;179(4):2313–7.

36. Maxwell PH, Curcio MJ. Host factors that control long terminal repeat retrotransposons in Saccharomyces cerevisiae: implications for regulation of mammalian retroviruses. Eukaryot Cell. 2007;6(7):1069–80.

37. Sankowski R, Strohl JJ, Huerta TS, Nasiri E, Mazzarello AN, D’Abramo C, et al. Endogenous retroviruses are associated with hippocampus-based memory impairment. Proc Natl Acad Sci U S A. 2019;116(51):25982–90.

38. Criscione SW, Zhang Y, Thompson W, Sedivy JM, Neretti N. Transcriptional landscape of repetitive elements in normal and cancer human cells. BMC Genomics. 2014;15:583.

39. Lee E, Iskow R, Yang L, Gokcumen O, Haseley P, Luquette LJ, 3rd, et al. Landscape of somatic retrotransposition in human cancers. Science. 2012;337(6097):967–71.

40. Lock FE, Rebollo R, Miceli-Royer K, Gagnier L, Kuah S, Babaian A, et al. Distinct isoform of FABP7 revealed by screening for retroelement-activated genes in diffuse large B-cell lymphoma. Proc Natl Acad Sci U S A. 2014;111(34):E3534–43.

41. Miki Y, Nishisho I, Horii A, Miyoshi Y, Utsunomiya J, Kinzler KW, et al. Disruption of the APC gene by a retrotransposal insertion of L1 sequence in a colon cancer. Cancer Res. 1992;52(3):643–5.

42. Rodriguez-Martin C, Cidre F, Fernandez-Teijeiro A, Gomez-Mariano G, de la Vega L, Ramos P, et al. Familial retinoblastoma due to intronic LINE-1 insertion causes aberrant and noncanonical mRNA splicing of the RB1 gene. J Hum Genet. 2016;61(5):463–6.

43. Scarfo I, Pellegrino E, Mereu E, Kwee I, Agnelli L, Bergaggio E, et al. Identification of a new subclass of ALK-negative ALCL expressing aberrant levels of ERBB4 transcripts. Blood. 2016;127(2):221–32.

44. Scott EC, Gardner EJ, Masood A, Chuang NT, Vertino PM, Devine SE. A hot L1 retrotransposon evades somatic repression and initiates human colorectal cancer. Genome Res. 2016;26(6):745–55.

45. Teugels E, De Brakeleer S, Goelen G, Lissens W, Sermijn E, De Greve J. De novo Alu element insertions targeted to a sequence common to the BRCA1 and BRCA2 genes. Hum Mutat. 2005;26(3):284.

46. Wiesner T, Lee W, Obenauf AC, Ran L, Murali R, Zhang QF, et al. Alternative transcription initiation leads to expression of a novel ALK isoform in cancer. Nature. 2015;526(7573):453–7.

47. Wolff EM, Byun HM, Han HF, Sharma S, Nichols PW, Siegmund KD, et al. Hypomethylation of a LINE-1 promoter activates an alternate transcript of the MET oncogene in bladders with cancer. PLoS Genet. 2010;6(4):e1000917.

48. Nguyen THM, Carreira PE, Sanchez-Luque FJ, Schauer SN, Fagg AC, Richardson SR, et al. L1 Retrotransposon Heterogeneity in Ovarian Tumor Cell Evolution. Cell Rep. 2018;23(13):3730–40.

49. Schauer SN, Carreira PE, Shukla R, Gerhardt DJ, Gerdes P, Sanchez-Luque FJ, et al. L1 retrotransposition is a common feature of mammalian hepatocarcinogenesis. Genome Res. 2018;28(5):639–53.

50. Shukla R, Upton KR, Munoz-Lopez M, Gerhardt DJ, Fisher ME, Nguyen T, et al. Endogenous retrotransposition activates oncogenic pathways in hepatocellular carcinoma. Cell. 2013;153(1):101–11.

51. Rodic N, Sharma R, Sharma R, Zampella J, Dai L, Taylor MS, et al. Long interspersed element-1 protein expression is a hallmark of many human cancers. Am J Pathol. 2014;184(5):1280–6.

52. Rodic N, Steranka JP, Makohon-Moore A, Moyer A, Shen P, Sharma R, et al. Retrotransposon insertions in the clonal evolution of pancreatic ductal adenocarcinoma. Nat Med. 2015;21(9):1060–4.

53. Treger RS, Pope SD, Kong Y, Tokuyama M, Taura M, Iwasaki A. The Lupus Susceptibility Locus Sgp3 Encodes the Suppressor of Endogenous Retrovirus Expression SNERV. Immunity. 2019;50(2):334–47 e9.

54. Wu Z, Mei X, Zhao D, Sun Y, Song J, Pan W, et al. DNA methylation modulates HERV-E expression in CD4+ T cells from systemic lupus erythematosus patients. J Dermatol Sci. 2015;77(2):110–6.

55. Neidhart M, Rethage J, Kuchen S, Kunzler P, Crowl RM, Billingham ME, et al. Retrotransposable L1 elements expressed in rheumatoid arthritis synovial tissue: association with genomic DNA hypomethylation and influence on gene expression. Arthritis Rheum. 2000;43(12):2634–47.

56. Arru G, Mameli G, Deiana GA, Rassu AL, Piredda R, Sechi E, et al. Humoral immunity response to human endogenous retroviruses K/W differentiates between amyotrophic lateral sclerosis and other neurological diseases. Eur J Neurol. 2018;25(8):1076–e84.

57. Douville R, Liu J, Rothstein J, Nath A. Identification of active loci of a human endogenous retrovirus in neurons of patients with amyotrophic lateral sclerosis. Ann Neurol. 2011;69(1):141–51.

58. Krug L, Chatterjee N, Borges-Monroy R, Hearn S, Liao WW, Morrill K, et al. Retrotransposon activation contributes to neurodegeneration in a Drosophila TDP-43 model of ALS. PLoS Genet. 2017;13(3):e1006635.

59. Li W, Jin Y, Prazak L, Hammell M, Dubnau J. Transposable elements in TDP-43-mediated neurodegenerative disorders. PLoS One. 2012;7(9):e44099.

60. Li W, Lee MH, Henderson L, Tyagi R, Bachani M, Steiner J, et al. Human endogenous retrovirus-K contributes to motor neuron disease. Sci Transl Med. 2015;7(307):307ra153.

61. Tam OH, Rozhkov NV, Shaw R, Kim D, Hubbard I, Fennessey S, et al. Postmortem Cortex Samples Identify Distinct Molecular Subtypes of ALS: Retrotransposon Activation, Oxidative Stress, and Activated Glia. Cell Rep. 2019;29(5):1164–77 e5.

62. Crow YJ, Rehwinkel J. Aicardi-Goutieres syndrome and related phenotypes: linking nucleic acid metabolism with autoimmunity. Hum Mol Genet. 2009;18(R2):R130–6.

63. Thomas CA, Tejwani L, Trujillo CA, Negraes PD, Herai RH, Mesci P, et al. Modeling of TREX1-Dependent Autoimmune Disease using Human Stem Cells Highlights L1 Accumulation as a Source of Neuroinflammation. Cell Stem Cell. 2017;21(3):319–31 e8.

64. Bollati V, Galimberti D, Pergoli L, Dalla Valle E, Barretta F, Cortini F, et al. DNA methylation in repetitive elements and Alzheimer disease. Brain Behav Immun. 2011;25(6):1078–83.

65. Guo C, Jeong HH, Hsieh YC, Klein HU, Bennett DA, De Jager PL, et al. Tau Activates Transposable Elements in Alzheimer’s Disease. Cell Rep. 2018;23(10):2874–80.

66. Protasova MS, Gusev FE, Grigorenko AP, Kuznetsova IL, Rogaev EI, Andreeva TV. Quantitative Analysis of L1-Retrotransposons in Alzheimer’s Disease and Aging. Biochemistry (Mosc). 2017;82(8):962–71.

67. Sun W, Samimi H, Gamez M, Zare H, Frost B. Pathogenic tau-induced piRNA depletion promotes neuronal death through transposable element dysregulation in neurodegenerative tauopathies. Nat Neurosci. 2018;21(8):1038–48.

68. Yan Z, Zhou Z, Wu Q, Chen ZB, Koo EH, Zhong S. Presymptomatic Increase of an Extracellular RNA in Blood Plasma Associates with the Development of Alzheimer’s Disease. Curr Biol. 2020;30(10):1771–82 e3.

69. Perron H, Bernard C, Bertrand JB, Lang AB, Popa I, Sanhadji K, et al. Endogenous retroviral genes, Herpesviruses and gender in Multiple Sclerosis. J Neurol Sci. 2009;286(1-2):65–72.

70. Perron H, Garson JA, Bedin F, Beseme F, Paranhos-Baccala G, Komurian-Pradel F, et al. Molecular identification of a novel retrovirus repeatedly isolated from patients with multiple sclerosis. The Collaborative Research Group on Multiple Sclerosis. Proc Natl Acad Sci U S A. 1997;94(14):7583–8.

71. Perron H, Suh M, Lalande B, Gratacap B, Laurent A, Stoebner P, et al. Herpes simplex virus ICP0 and ICP4 immediate early proteins strongly enhance expression of a retrovirus harboured by a leptomeningeal cell line from a patient with multiple sclerosis. J Gen Virol. 1993;74 (Pt 1):65–72.

72. Tan H, Qurashi A, Poidevin M, Nelson DL, Li H, Jin P. Retrotransposon activation contributes to fragile X premutation rCGG-mediated neurodegeneration. Hum Mol Genet. 2012;21(1):57–65.

73. Gemenetzi M, Lotery AJ. The role of epigenetics in age-related macular degeneration. Eye (Lond). 2014;28(12):1407–17.

74. Zhao B, Wu Q, Ye AY, Guo J, Zheng X, Yang X, et al. Somatic LINE-1 retrotransposition in cortical neurons and non-brain tissues of Rett patients and healthy individuals. PLoS Genet. 2019;15(4):e1008043.

75. Bourque G, Burns KH, Gehring M, Gorbunova V, Seluanov A, Hammell M, et al. Ten things you should know about transposable elements. Genome Biol. 2018;19(1):199.

76. Wicker T, Sabot F, Hua-Van A, Bennetzen JL, Capy P, Chalhoub B, et al. A unified classification system for eukaryotic transposable elements. Nat Rev Genet. 2007;8(12):973–82.

77. Boeke JD, Garfinkel DJ, Styles CA, Fink GR. Ty elements transpose through an RNA intermediate. Cell. 1985;40(3):491–500.

78. Ashley J, Cordy B, Lucia D, Fradkin LG, Budnik V, Thomson T. Retrovirus-like Gag Protein Arc1 Binds RNA and Traffics across Synaptic Boutons. Cell. 2018;172(1-2):262–74 e11.

79. Kawamura Y, Sanchez Calle A, Yamamoto Y, Sato TA, Ochiya T. Extracellular vesicles mediate the horizontal transfer of an active LINE-1 retrotransposon. J Extracell Vesicles. 2019;8(1):1643214.

80. Pastuzyn ED, Day CE, Kearns RB, Kyrke-Smith M, Taibi AV, McCormick J, et al. The Neuronal Gene Arc Encodes a Repurposed Retrotransposon Gag Protein that Mediates Intercellular RNA Transfer. Cell. 2018;173(1):275.

81. Ribet D, Harper F, Dupressoir A, Dewannieux M, Pierron G, Heidmann T. An infectious progenitor for the murine IAP retrotransposon: emergence of an intracellular genetic parasite from an ancient retrovirus. Genome Res. 2008;18(4):597–609.

82. Burke WD, Malik HS, Rich SM, Eickbush TH. Ancient lineages of non-LTR retrotransposons in the primitive eukaryote, Giardia lamblia. Mol Biol Evol. 2002;19(5):619–30.

83. Eickbush DG, Eickbush TH. Vertical transmission of the retrotransposable elements R1 and R2 during the evolution of the Drosophila melanogaster species subgroup. Genetics. 1995;139(2):671–84.

84. Malik HS, Burke WD, Eickbush TH. The age and evolution of non-LTR retrotransposable elements. Mol Biol Evol. 1999;16(6):793–805.

85. Malik HS, Henikoff S, Eickbush TH. Poised for contagion: evolutionary origins of the infectious abilities of invertebrate retroviruses. Genome Res. 2000;10(9):1307–18.

86. Jones BC, Wood JG, Chang C, Tam AD, Franklin MJ, Siegel ER, et al. A somatic piRNA pathway in the Drosophila fat body ensures metabolic homeostasis and normal lifespan. Nat Commun. 2016;7:13856.

87. Sousa-Victor P, Ayyaz A, Hayashi R, Qi Y, Madden DT, Lunyak VV, et al. Piwi Is Required to Limit Exhaustion of Aging Somatic Stem Cells. Cell Rep. 2017;20(11):2527–37.

88. Zhou F, Li M, Wei Y, Lin K, Lu Y, Shen J, et al. Activation of HERV-K Env protein is essential for tumorigenesis and metastasis of breast cancer cells. Oncotarget. 2016;7(51):84093–117.

89. Chan SM, Sapir T, Park SS, Rual JF, Contreras-Galindo R, Reiner O, et al. The HERV-K accessory protein Np9 controls viability and migration of teratocarcinoma cells. PLoS One. 2019;14(2):e0212970.

90. Gonzalez-Cao M, Iduma P, Karachaliou N, Santarpia M, Blanco J, Rosell R. Human endogenous retroviruses and cancer. Cancer Biol Med. 2016;13(4):483–8.

91. Kim A, Terzian C, Santamaria P, Pelisson A, Purd’homme N, Bucheton A. Retroviruses in invertebrates: the gypsy retrotransposon is apparently an infectious retrovirus of Drosophila melanogaster. Proc Natl Acad Sci U S A. 1994;91(4):1285–9.

92. Marlor RL, Parkhurst SM, Corces VG. The Drosophila melanogaster gypsy transposable element encodes putative gene products homologous to retroviral proteins. Mol Cell Biol. 1986;6(4):1129–34.

93. Pelisson A, Song SU, Prud’homme N, Smith PA, Bucheton A, Corces VG. Gypsy transposition correlates with the production of a retroviral envelope-like protein under the tissue-specific control of the Drosophila flamenco gene. EMBO J. 1994;13(18):4401–11.

94. Song SU, Gerasimova T, Kurkulos M, Boeke JD, Corces VG. An env-like protein encoded by a Drosophila retroelement: evidence that gypsy is an infectious retrovirus. Genes Dev. 1994;8(17):2046–57.

95. Ribet D, Harper F, Esnault C, Pierron G, Heidmann T. The GLN family of murine endogenous retroviruses contains an element competent for infectious viral particle formation. J Virol. 2008;82(9):4413–9.

96. Dewannieux M, Harper F, Richaud A, Letzelter C, Ribet D, Pierron G, et al. Identification of an infectious progenitor for the multiple-copy HERV-K human endogenous retroelements. Genome Res. 2006;16(12):1548–56.

97. Robinson-McCarthy LR, McCarthy KR, Raaben M, Piccinotti S, Nieuwenhuis J, Stubbs SH, et al. Reconstruction of the cell entry pathway of an extinct virus. PLoS Pathog. 2018;14(8):e1007123.

98. Szymczak AL, Workman CJ, Wang Y, Vignali KM, Dilioglou S, Vanin EF, et al. Correction of multi-gene deficiency in vivo using a single ‘self-cleaving’ 2A peptide-based retroviral vector. Nat Biotechnol. 2004;22(5):589–94.

99. Capy P, Langin T, Higuet D, Maurer P, Bazin C. Do the integrases of LTR-retrotransposons and class II element transposases have a common ancestor? Genetica. 1997;100(1-3):63–72.

100. Chalvet F, Teysset L, Terzian C, Prud’homme N, Santamaria P, Bucheton A, et al. Proviral amplification of the Gypsy endogenous retrovirus of Drosophila melanogaster involves env-independent invasion of the female germline. EMBO J. 1999;18(9):2659–69.

101. Syomin BV, Fedorova LI, Surkov SA, Ilyin YV. The endogenous Drosophila melanogaster retrovirus gypsy can propagate in Drosophila hydei cells. Mol Gen Genet. 2001;264(5):588–94.

102. Teysset L, Burns JC, Shike H, Sullivan BL, Bucheton A, Terzian C. A Moloney murine leukemia virus-based retroviral vector pseudotyped by the insect retroviral gypsy envelope can infect Drosophila cells. J Virol. 1998;72(1):853–6.

103. Bodea GO, McKelvey EGZ, Faulkner GJ. Retrotransposon-induced mosaicism in the neural genome. Open Biol. 2018;8(7).

104. Upton KR, Gerhardt DJ, Jesuadian JS, Richardson SR, Sanchez-Luque FJ, Bodea GO, et al. Ubiquitous L1 mosaicism in hippocampal neurons. Cell. 2015;161(2):228–39.

105. Maxwell PH. What might retrotransposons teach us about aging? Curr Genet. 2016;62(2):277–82.

106. Savva YA, Jepson JE, Chang YJ, Whitaker R, Jones BC, St Laurent G, et al. RNA editing regulates transposon-mediated heterochromatic gene silencing. Nat Commun. 2013;4:2745.

107. Simon M, Van Meter M, Ablaeva J, Ke Z, Gonzalez RS, Taguchi T, et al. LINE1 Derepression in Aged Wild-Type and SIRT6-Deficient Mice Drives Inflammation. Cell Metab. 2019;29(4):871–85 e5.

108. Blaudin de The FX, Rekaik H, Peze-Heidsieck E, Massiani-Beaudoin O, Joshi RL, Fuchs J, et al. Engrailed homeoprotein blocks degeneration in adult dopaminergic neurons through LINE-1 repression. EMBO J. 2018;37(15).

109. Kaneko H, Dridi S, Tarallo V, Gelfand BD, Fowler BJ, Cho WG, et al. DICER1 deficit induces Alu RNA toxicity in age-related macular degeneration. Nature. 2011;471(7338):325–30.

110. Tarallo V, Hirano Y, Gelfand BD, Dridi S, Kerur N, Kim Y, et al. DICER1 loss and Alu RNA induce age-related macular degeneration via the NLRP3 inflammasome and MyD88. Cell. 2012;149(4):847–59.

111. Carreira PE, Ewing AD, Li G, Schauer SN, Upton KR, Fagg AC, et al. Evidence for L1-associated DNA rearrangements and negligible L1 retrotransposition in glioblastoma multiforme. Mob DNA. 2016;7:21.

112. Doucet-O’Hare TT, Rodic N, Sharma R, Darbari I, Abril G, Choi JA, et al. LINE-1 expression and retrotransposition in Barrett’s esophagus and esophageal carcinoma. Proc Natl Acad Sci U S A. 2015;112(35):E4894–900.

113. Doucet-O’Hare TT, Sharma R, Rodic N, Anders RA, Burns KH, Kazazian HH, Jr. Somatically Acquired LINE-1 Insertions in Normal Esophagus Undergo Clonal Expansion in Esophageal Squamous Cell Carcinoma. Hum Mutat. 2016;37(9):942–54.

114. Solyom S, Ewing AD, Rahrmann EP, Doucet T, Nelson HH, Burns MB, et al. Extensive somatic L1 retrotransposition in colorectal tumors. Genome Res. 2012;22(12):2328–38.

115. Tang Z, Steranka JP, Ma S, Grivainis M, Rodic N, Huang CR, et al. Human transposon insertion profiling: Analysis, visualization and identification of somatic LINE-1 insertions in ovarian cancer. Proc Natl Acad Sci U S A. 2017;114(5):E733–E40.

116. Wylie A, Jones AE, D’Brot A, Lu WJ, Kurtz P, Moran JV, et al. p53 genes function to restrain mobile elements. Genes Dev. 2016;30(1):64–77.

117. Yang N, Kazazian HH, Jr. L1 retrotransposition is suppressed by endogenously encoded small interfering RNAs in human cultured cells. Nat Struct Mol Biol. 2006;13(9):763–71.

118. Carreira PE, Richardson SR, Faulkner GJ. L1 retrotransposons, cancer stem cells and oncogenesis. FEBS J. 2014;281(1):63–73.

