## supplementary figure 1 for "Intercellular viral spread and intracellular transposition of *Drosophila* gypsy"

A

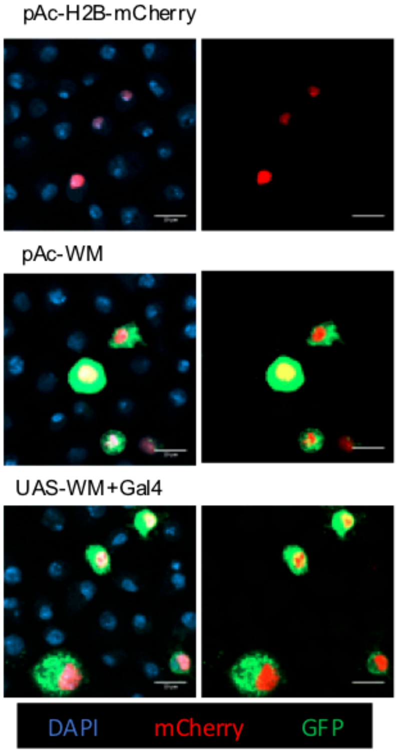


B

| Plasmid (s) Transfected | Cells Counted | Cells Labeled (%) |
|---|---|---|
| pAc-H2B-mCherry | 9148 | 365 (3.98) |
| pAc-WM | 9186 | 881 (9.59) |
| UAS-WM no Gal4 | 9079 | 0 |
| UAS-WM Tub Gal4 | 9173 | 947 (10.3) |
