## supplementary figure 2 for "Intercellular viral spread and intracellular transposition of *Drosophila* gypsy"

A

Gypsy-Env Nucleotide Sequence:

atGTTCACCCTCATGATGTTCATACCCTTGGTAGTAGCGAATGCTCGGATCACCGACTTTTCGCATGCCAACTACAT
TCCTGTGTTAGATGGGGATGTGCTGGTGTTTGAACAGCGTGACCTCTTGAAACATTCGAGTAACCTTTCCGAGTACG
CTAGTATGATAGATGAAACACAGAAACTGTCCGAGTCCTTTCCCCACTCACATATGCGTAAGTTGCTAGAGGTCGAT
ACTGACCATCTTAGAACCTTGTTGTCCGTTCTCAAAGTCCACCATAGGATAGCTAGGAGTCTAGATTTCTTAGGTAC
AGCCTTAAAGGTTGTGGCGGGTACTCCCGATGCCACGGACCTCTTTAAAATTAAGATCACAGAGGCCCAACTAGTAG
AATCTAATTCCAGGCAGATAGCTATAAACTCCGAAACCCAGAAACAGATAAATAAGTTAACTGACACCATCAATAAG
GTGATCAATGCCCGTAAAGGCGACTTGGTTGACACTCCACACTTATATGAAGCACTACTAGCAAGAAATAGGATGCT
GTCTACAGAAATTCAAAATTTAATTCTCACTATTACTTTGGTCAAATCAAACATTATAAATCCCACAATTCTTGATC
ATGCCGACTTGAAGCCTCTTGTAGAACAGGATACCCCAATTGTCAGCTTAATAGAAGCATCTAAGATCAGGGTCCTC
CAGTCCGAGAATAGCATTCATATTTTAATTGCCTATCCTAGAGTCAAGTTCAGTTGCAAGAAAGTCGCCGTCTACCC
TGTATCTCACCAACACACCATCTTGCGCCTCGACGAAGACACTTTGGCCGAATGCGAACATGACACCTTTGCGGTCA
CCGGATGCACAGACACCACACACTTCACGTTCTGCGAGCGGTCTCGGCGCGAAACTTGCGTGCGCTCACTCCATGCT
GGAAACGCTGCTCAATGCCACACTCAACCCAGCCACTTGCGAGAAATAAACCCCGTAGATGATGGCGTTGTGATTAT
CAACGAAGCCGCAGCTCACGTTAGCACTGATGGCAGCCCCGAAACACTGATAGAGGGAACCTACCTGGTAACCTTCG
AGCGAACGGCAACCATCAACGGCTCTGAATTCGTAAATCTAAGGAAAACACTAAGCAAGCAGCCAGGCATCGTGCGT
TCACCACTACTTAACATCGTCGGCCACGACCCTGTGCTCAGTATACCTCTGCTACACCGGATGAGTAACGAAAACCT
ACATTCCATCCAAAACCTTATGGATGACGTGGAATCTGAAGGCTCGCCCAGACTCTGGTTCGTGGCTGGTGTGGTCC
TAAACTTCGGCTTGATTGGCTCTCTCGCCCTTTATCTGGCATTAAGGAGAAGACGAGCCTCTAGGGAGATACAGCGC
ACCATCGATACTTTCAACATGACCGAGGACGGTCATAAACTTGAGGGGGGAGTAGTTAACAACTAA

B

Gypsy-Env Amino Acid Sequence:

MFTLMMFIPLVVANARITDFSHANYIPVLDGDVLVFEQRDLLKHSSNLSEYASMIDETQKLSESFPHSHMRKLLEVD
TDHLRTLLSVLKVHHRIARSLDFLGTALKVVAGTPDATDLFKIKITEAQLVESNSRQIAINSETQKQINKLTDTINK
VINARKGDLVDTPHLYEALLARNRMLSTEIQNLILTITLVKSNIINPTILDHADLKPLVEQDTPIVSLIEASKIRVL
QSENSIHILIAYPRVKFSCKKVAVYPVSHQHTILRLDEDTLAECEHDTFAVTGCTDTTHFTFCERSRRETCVRSLHA
GNAAQCHTQPSHLREINPVDDGVVIINEAAAHVSTDGSPETLIEGTYLVTFERTATINGSEFVNLRKTLSKQPGIVR
SPLLNIVGHDPVLSIPLLHRMSNENLHSIQNLMDDVESEGSPRLWFVAGVVLNFGLIGSLALYLALRRRRASREIQR
TIDTFNMTEDGHKLEGGVVNN-

C

```
CLUSTAL O(1.2.4) multiple sequence alignment


Env             MFTLMMFIPLVVANARITDFSHANYIPVLDGDVLVFEQRDLLKHSSNLSEYASMIDETQK
Env-deletion    MFTLMMFIPLVVANARITDFSHANYIPVLDGDVLVFEQRDLLKHSSNLSEYASMIDETQK
                ************************************************************

Env             LSESFPHSHMRKLLEVDTDHLRTLLSVLKVHHRIARSLDFLGTALKVVAGTPDATDLFKI
Env-deletion    LSESFPHSHMRKLLEVDTDHLRTLLSVLKVHHRIGV*-----------------------
                **********************************.

Env             KITEAQLVESNSRQIAINSETQKQINKLTDTINKVINARKGDLVDTPHLYEALLARNRML
Env-deletion    ------------------------------------------------------------

Env             STEIQNLILTITLVKSNIINPTILDHADLKPLVEQDTPIVSLIEASKIRVLQSENSIHIL
Env-deletion    ------------------------------------------------------------

Env             IAYPRVKFSCKKVAVYPVSHQHTILRLDEDTLAECEHDTFAVTGCTDTTHFTFCERSRRE
Env-deletion    ------------------------------------------------------------

Env             TCVRSLHAGNAAQCHTQPSHLREINPVDDGVVIINEAAAHVSTDGSPETLIEGTYLVTFE
Env-deletion    ------------------------------------------------------------

Env             RTATINGSEFVNLRKTLSKQPGIVRSPLLNIVGHDPVLSIPLLHRMSNENLHSIQNLMDD
Env-deletion    ------------------------------------------------------------

Env             VESEGSPRLWFVAGVVLNFGLIGSLALYLALRRRRASREIQRTIDTFNMTEDGHKLEGGV
Env-deletion    ------------------------------------------------------------

Env             VNN*
Env-deletion    ----
```
